# A gene-centric analysis of copy number variation reveals tumour suppressor modulation of cancer risk and novel associations with common diseases

**DOI:** 10.64898/2026.10.06.757170

**Authors:** Sophie Matthews, Cathal Seoighe, Gregory L. Wheeler

## Abstract

Copy number variants (CNVs) are a significant source of genomic variation in humans. They can span large genomic regions that encompass whole genes and their flanking regions, having the potential to alter the number of functional copies in an individual. Typically, CNVs are analysed individually or combined into regions without regard to the genes they span. Here, we employ a gene-centric approach to link changes in whole-gene copy number to phenotype. We first examined the relationship between CNVs spanning whole genes and gene expression using data from GTEx and found that predicted effects on gene expression are often realised. Applying this method, we identified 34 gene deletions associated with cancer incidence, including several novel candidate associations. Most notably, we discovered that gene deletions in the 13q14 region are linked to increased cancer risk, a novel extension of a known somatic association to the germline, which we did not find using region-based methods. Statistical associations were found between whole-gene loss of tumour suppressor genes and overall cancer risk, as hypothesized. It was additionally found that the total number of whole-gene losses was strongly associated with increased cancer risk, even when known cancer-related genes are not included in the count. Finally, we extended these methods to non-cancer phenotypes, uncovering significant associations with common diseases, including asthma and heart attack.

## Introduction

Copy number variants (CNVs) are regions of the genome that vary in copy number relative to the reference genome, comprising both duplications and deletions of DNA. While many CNVs are tolerated without apparent effect (often being small, intergenic or encompassing genes that can tolerate copy number changes), others are strongly associated with diverse human diseases (Y. R. Li et al., 2020; Malhotra & Sebat, 2012; Weischenfeldt et al., 2013). CNVs often span entire genes, altering their copy number between individuals; however, they are often analysed as large, multi-gene regions. This approach can obscure the specific role of individual genes within the interval, making it difficult to distinguish which gene(s) are driving the phenotypic effect.

To enable genome-wide association testing, CNVs are typically aggregated into non-overlapping CNV regions (CNVRs) based on the overlap of calls across individuals. However, there is no standardised method for this aggregation. Common strategies include density-based approaches (Kim et al., 2012), which merge adjacent CNVs based on the proportion of individuals sharing an identical segment, and reciprocal overlap methods (Conrad et al., 2010; da Silva et al., 2019), which require two CNVs to overlap each other by a specified fraction of their lengths. The resulting set of CNVRs are then dependent on the choice of aggregation method and its parameters (e.g. density or overlap thresholds), which can influence the associations detected. To overcome this limitation for gene-based hypotheses, a gene-centric framework is needed, where individual genes, rather than large genomic windows, are the primary focus of analysis. Previous studies have successfully employed gene-focused methods, associating impacted gene clusters from large CNVs with disease prevalence and protein biomarker levels (Zou et al., 2026).

Studies have also considered the effects of smaller copy-impacting changes such as structural variants (Bai et al., 2026). This study bridges both areas, covering both megabase-scale CNV events impacting dozens of genes as well as focal CNVs of single exons.

The need for a gene-level view is particularly critical in the case of dosage sensitive genes, where gains or losses in copy number can have significant, and often pathogenic, phenotypic effects (Rice & McLysaght, 2017). A reduction in gene copies can result in haploinsufficiency, whereby decreased copy number reduces gene product and impairs function (Fisher & Scambler, 1994). In addition, duplications of dosage-sensitive genes can also contribute to the pathogenicity of CNVs by disrupting normal gene function (Lupski et al., 1991). However, not all dosage sensitivity is harmful; for example, an increased copy number of *AMY1 is* associated with protection against obesity and metabolic disorders and is proposed to improve glucose metabolism and insulin sensitivity (Greenhill, 2014; León-Mimila et al., 2018).

The “two-hit” model of cancer causation is another example whereby a change in gene copy number could be advantageous. The two-hit model proposes that, for a tumour suppressor gene (TSG) to contribute to cancer, both copies of the gene must be inactivated, whether this be through mutations or deletions (Knudson, 1971). A gain in copy number for a TSG would require additional loss-of-function events to fully inactivate the gene, potentially lowering cancer risk. Therefore, we hypothesised that an increase in TSG copy number could confer protection against cancer, whereas gene deletions may have the opposite effect. Deletions would leave only one functional copy, requiring just a single additional “hit” to inactivate it, thereby increasing an individual’s cancer risk.

In contrast to TSGs, cancer-causing mutations in oncogenes are typically dominant, such that a single mutation in one copy of the gene is sufficient to drive cancer progression. Therefore, we hypothesised that, in addition to the direct contribution via increased activity of the oncogene, an increase in the number of functional copies of an oncogene may increase the chance of a gene acquiring an activating mutation. On the other hand, by reducing the driver mutation target size, a deletion may reduce the probability that the oncogene acquires an activating mutation. The X chromosome provides a natural test case of this hypothesis: females have two copies of most X-linked genes, whereas males only have one, enabling us to examine how gene copy number influences the accumulation of somatic mutations. Previous work has associated interspecific variation in copy-number of both TSGs and oncogenes with prevalence of cancer (Matthews, 2026; Matthews et al., 2025), suggesting selection and functional constraint in genes of these types.

In this study, we implement a gene-centric framework to investigate the phenotypic consequences of whole-gene copy number variation. We first assessed the dosage effects of gene copy number on gene expression using data from the GTEx consortium. Motivated by the hypothesis that additional copies of oncogenes could elevate cancer risk by increasing opportunities for activating mutations, we then compared mutation rates of X-linked genes between males and females. Consistent with expectations, genes lacking a Y-linked homolog showed approximately double the mutation rate in females compared to males. We next tested the hypothesis that variable copy number of TSGs and oncogenes could influence cancer risk in an individual, finding evidence that supports this hypothesis. Finally, we expanded this gene-centric approach to perform genome-wide association tests between the copy number of every gene and risk for cancer and other common diseases, thereby identifying specific genes whose dosage may contribute to disease susceptibility.

## Methods

### CNV detection and expression analysis

To assess the impact of whole-gene copy number variation on expression levels, we called CNVs from whole-genome sequencing (WGS) CRAM files for 908 individuals from the GTEx version 8 release (GTEx Consortium, 2020). CNVs were called from these files using CNVpytor (version 1.2.1, (Suvakov et al., 2021), a python extension of its predecessor CNVnator (Abyzov et al., 2011), which applies a read-depth approach to CNV detection. CNV calling was performed using a bin size of 1000 bp, consistent with the recommended minimal bin size outlined by Suvakov et al. (2021). CNV calls were filtered to exclude those with more than 50% of zero quality (q0) mapped reads as described by Abyzov et al. (2011).

Genes with a variable copy number were identified from the CNV calls based on gene annotation of GRCh38 obtained from Gencode (version 41, (Frankish et al., 2023). Genes with a variable copy number were defined as whole genes located within CNV boundaries. To increase the likelihood that CNVs encompassed entire genes, including promoter regions outside annotated boundaries, we expanded the gene boundaries by 1 kb on either side.

Whole blood gene expression data was downloaded from the GTEx consortium version 8 release (GTEx Consortium, 2020). A total of 680 individuals had both gene expression and WGS data available. To include only expressed genes, expression data was filtered for genes with a median transcript per million (TPM) value of greater than 0.5.

Spearman’s correlation coefficient was used to assess the association between gene copy number and gene expression. Genes included in the analysis were filtered to retain only those with variable copy numbers in at least 5% of individuals. Correlation tests were conducted for all variable copy numbers and were subsequently repeated separately for deletions and duplications. We tested for enrichment of immune genes among genes showing a significant correlation between copy number and expression using Fisher’s exact test. The background set included all genes tested for correlation (i.e. those with observed duplications or deletions), controlling for higher CNV frequency among immune genes. Immune-related genes were identified using annotations from InnateDB (version 5.4, (Breuer et al., 2013).

### X-linked mutation analysis

To provide proof-of-principle for our hypothesis that an increased copy number could increase the chance of an oncogene acquiring an activating mutation, we evaluated whether mutational burden scales with copy number, using the X chromosome as a natural test case. Somatic mutation data was obtained from the COSMIC gene census (version 101, (Sondka et al., 2018). Using data from (Balaton et al., 2015), we identified genes on the X chromosome present within the pseudoautosomal region (PAR), therefore having a copy on the Y chromosome and retaining a copy number of two in both males and females. For each sex, we aggregated the total number of mutations in the database and calculated the proportion of mutations on genes that were uniquely X-linked i.e. not in the PAR. We performed a chi-squared test to determine if there was a significant difference in the proportion of mutations on these genes between males and females. This group of genes was then further separated into genes that are inactivated and those that escape inactivation. We also calculated the proportion of total mutations in the PAR and subsequently performed a chi-squared test to determine if mutations in the PAR differed between sexes.

### Gene copy number and cancer risk

We next tested whether variation in copy number of TSGs or oncogenes is associated with cancer incidence, under the hypothesis that duplications of TSGs may be protective, while deletions may increase risk, with the opposite being true for oncogenes. CNVs were called for 488,377 participants from the UK Biobank (http://www.ukbiobank.ac.uk) using PennCNV v1.0.5 (K. Wang et al., 2007), based on log R ratio and B allele frequency data derived from the UK Biobank genotype dataset (UKB category 263). Genetic data acquisition and quality control was performed by the UK Biobank and have been previously described (Bycroft et al., 2018). CNV calling was conducted using PennCNV’s general mode, which applies likelihood-based testing to infer discrete copy-number states (CN = 0, 1, 2, 3, 4) at each marker based on log-normalised signal intensity and allele balance in individual samples. The HMM model was initialised with conditions optimised for Affymetrix arrays (affygw6.hmm), as provided in the PennCNV resources. For each individual, gene copy number was inferred by identifying genes spanned by a CNV, including an additional 1 kb flanking region on either side of the gene, as previously described. The positions of markers in the UK Biobank data are reported in GRCh37 coordinates. To maintain consistency with this assembly, gene coordinates were obtained for GRCh37 from Ensembl Biomart (version 75;(Harrison et al., 2024). Partial-gene CNV events impacting a portion of gene coding sequence (CDS) were also identified. Coding sequences were inferred from each gene’s canonical transcript, identified from the GENCODE GRCh37 genome annotation (Frankish et al., 2023). To select a single representative transcript per gene, we prioritised transcripts with an APPRIS principal 1 annotation (Rodriguez et al., 2022). For genes lacking an APPRIS principal 1 annotation, we selected the highest available APPRIS principal level. Whole-gene and partial-gene CDS alterations were treated as mutually exclusive; in cases where both a whole-gene and partial-gene CNV of the same gene and type were detected in an individual, the whole-gene CNV was considered to take precedence. Participants found to have over 500 total CNVs (roughly corresponding to the 99.5th percentile) were considered to represent likely data quality issues or phenomena not representative of the broader population and were excluded from analyses.

Using the COSMIC gene census of cancer genes (Sondka et al., 2018), we identified TSGs and oncogenes within our dataset of genes with a variable copy number. We conducted logistic regression analyses to evaluate whether cancer incidence was associated with duplications or deletions of each gene class. Logistic regression analyses were carried out separately for deletions and duplications using the binomial test family, and accounted for covariates: age, sex, BMI, population structure (principal components 1–4), batch, and smoking status, with a binary predictor variable indicating presence or absence of any event of interest in the gene class of interest. Analyses were conducted for the full set of oncogenes and TSGs, and then repeated for a subject of these genes with known cancer driver variants as characterised by (Bailey et al., 2018) and repeated the analysis for these subsets. To exclude the confounding effects of artifactual CNVs and those common in the population, analyses were performed excluding CNVs found in over 1000 participants. . Statistical power for binary predictors was calculated using Hsieh’s formula (Hsieh, 1989).

To assess the impact of total CNV burden, logistic regression using numeric predictors was also conducted. Cancer incidence was compared to participants’ total numbers of whole-gene gains and losses as well as partial-gene gains and losses, again using the binomial test family and accounting for covariates: age, sex, BMI, population structure, batch, and smoking status.

### Genome-wide association tests

Finally, we applied our gene-centric framework on a genome-wide scale to explore associations between the copy number of every protein-coding gene and disease incidence. Using gene copy number data from the UKB cohort, logistic regression analyses were conducted to assess whether the deletion or duplication of any genes influenced cancer incidence. The logistic regression models included the following covariates: age, sex, population structure (principal components 1–4), batch, BMI and smoking status. We then separated out the three most common cancer types in the UKB cohort (skin, breast, and prostate cancer) and performed the same analyses to assess associations between gene copy number and the incidence of these specific cancer types. Subsequently, we expanded this approach to analyse additional phenotypes: angina, asthma, diabetes, deep vein thrombosis (DVT), heart attack, pulmonary embolism and stroke.

Additionally, we tested for associations between deletions that disrupt protein-coding genes, defined as deletions overlapping the canonical transcript as described above, and the same phenotypes.

To compare our gene-centric association study to standard CNV-based approaches, we conducted genome-wide association tests between cancer incidence and CNVs. CNVs were first aggregated into CNV regions (CNVRs; genomic intervals that define the minimal common region of overlapping CNVs across individuals). We employed two aggregation strategies for comparison to our gene-centric approach. Firstly, we employed a density-based procedure from CNVRuler (Kim et al., 2012), which merges adjacent CNVs based on a density threshold. This method defines a CNVR when a specified minimum proportion of individuals with overlapping CNVs in a region share an identical segment. We applied this using both a standard threshold (0.1, requiring 10% overlap) and a more sensitive threshold (0.05) to capture a broader set of regions. We then employed a reciprocal overlap approach, implemented in CNVRanger (da Silva et al., 2019). This method aggregates two CNVs if they reciprocally overlap each other by a minimum proportion of their respective lengths. We used the default reciprocal overlap threshold of 0.5.

For each of these methods, we performed logistic regression analyses to test for associations between CNVRs and cancer incidence, using the same covariates as in the gene-centric framework (age, sex, population structure [PC1–PC4], batch, BMI, and smoking status). This allowed direct comparison between gene-level copy number associations and CNV-based associations across the genome.

## Results

### Copy-number Distributions

A total of 408,926 unique CNVs were detected, with a CNV considered to be unique if either the start or end breakpoint differed from another, with gains and losses considered separately. Sizes ranged from as small as 20 bp to as large as 31.9 megabases, with a median size of 94.5 kilobases (Supplementary figure 1).

While the frequency of CNVs differed by orders of magnitude across genomic bins, overall the genome was well-covered by both gains and losses (Figure 1A). At the scale of the 2.5 megabase bins considered, most bins possessed hundreds of CNVs across the cohort. The few large regions that showed aberrant patterns were known low-complexity regions as in chromosomes 1 and 9; however, as these regions lack gene content, they are not expected to impact gene-centric analyses.

**Figure 1.**
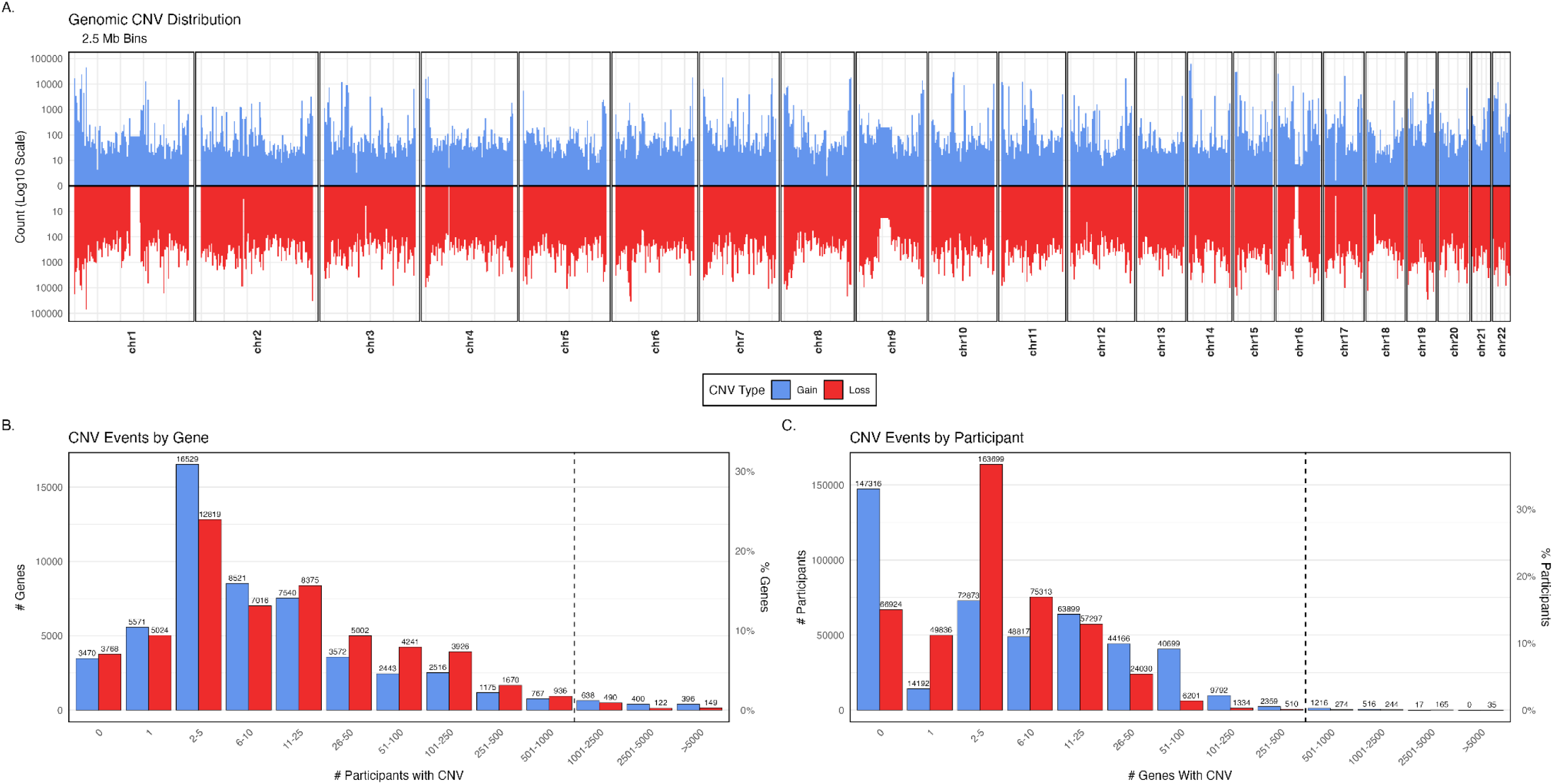
Plots showing distribution of copy-number variants – gains (blue) or losses (red) across the genome (**A**), genes (**B**), and participants (**C**). **1A.** Genomic CNV distribution. Positions plotted are per-chromosome, with each bar representing a 2.5 megabase genomic bin. Counts are log10 scaled and show the total of participants possessing a gain (upwards) or loss (downwards) impacting any part of the bin. **1B.** Rate of CNVs by genes. Bins are categorical and not to scale; dashed line indicates cutoff point for analyses which exclude common CNVs. **1C.** Rate of CNVs by individual participants. Bins categorical; dashed line indicates cutoff for individuals excluded from cancer risk analysis due to abnormally high CNV burden.

At the scale of individual genes (Figure 1B), the majority of the 53,538 genes tested were spanned by CNVs in only a small number of participants (median: 6 individuals with whole-gene gains and 9 with whole-gene losses). A small minority were not observed to have any gene-spanning CNVs at all (6.48% of genes had no whole-gene gains in any individual and 7.03% had no whole-gene losses). Conversely, a smaller minority (gains: 2.68%; losses: 1.42%) had gains or losses observed in over 1000 participants; as it is unclear if these high-frequency events indicate shared ancestry or recurrent statistical artifacts, these events were excluded from analyses.

The burden of gene-spanning CNVs per individual was also assessed (Figure 1C), with the median individual found to have 9 genes with an increase or decrease in gene copy number as a result of a CNV (gains: 5, losses: 4). A substantial portion of the cohort possessed no whole-gene gains (33.0%), with fewer possessing no whole-gene losses (15.0%). The cohort was observed to have a “long tail” of individuals with very high CNV burdens: while an individual with over 250 genes impacted with gains or over 70 with losses falls into the 99th percentile of per-category CNV burden, individuals were present with much greater values (max: 3694 gains, 7104 losses). Such extreme values seem biologically implausible and may result from data quality issues; therefore, we excluded individuals with more than 500 whole-gene gains or losses from our subsequent analyses (roughly corresponding to the 99.5th percentile).

### Impact of gene copy number on expression

Among the 680 GTEx individuals with matched WGS and gene expression data, 76 genes had deletions and 38 genes had duplications of protein-coding genes that were shared with more than 5% of the individuals. Of these, 50/76 deletions (*Supplementary table 1*) and 29/38 duplications (*Supplementary table 2*) showed significant correlations with gene expression (*p* < 0.05). The majority followed the expected dosage pattern: 42 deletions were associated with reduced expression and 23 duplications with increased expression.

Among genes with observed copy number variation, immune genes were significantly more likely than non-immune genes to show significant associations between copy number and expression (Fisher’s test: deletions; *p =* 0.0037, duplications; *p =* 2.2e-08). Immune genes significantly associated with expression included immunoglobulin genes, T-cell receptor genes, and human leukocyte antigen (HLA) genes. Genes exhibiting reduced expression despite increased copy number were exclusively immune-related, namely T-cell receptor genes.

### Sex differences in the X chromosome mutation burden

As expected, the proportion of mutations affecting genes located exclusively on the X chromosome (i.e. not in the PAR) was significantly higher in females than in males (4.5% in females, compared to 2.4% in males; *p* < 2.2e-16; *Supplementary figure 2A*). In contrast, the proportion of total mutations located in PAR genes did not differ significantly between sexes (females = 0.071%, males = 0.076%, *p =* 0.54; *Supplementary figure 2B*). Further separation of X chromosome-exclusive genes into those that escape inactivation and those that are inactivated, revealed that both groups harbour a higher proportion of total mutations in females than males, though the effect appeared to be stronger in the X-inactivated genes (3.8% in female compared to 1.9% in male for inactivated genes and 0.71% compared to 0.55% in genes that escape inactivation).

### Assessing the impact of gene copy number on cancer risk

As hypothesized, whole-gene loss of TSGs with known drivers had a strong positive effect on cancer risk (Table 1, p = 0.033, OR: 1.098); whole-gene loss of TSGs overall was found to be marginal (p = 0.084, OR: 1.076), likely due to the effect of driver TSGs, which made up the vast majority of all TSGs affected. One additional marginal effect was detected (0.10 > α > 0.05), that of whole-gene losses on oncogenes with known drivers; however, this effect was in the opposite direction of that hypothesized. These losses were weakly supported to increase cancer risk, in opposition to the protective effect predicted by the reduced target area hypothesis (p = 0.064, OR: 1.076). All non-significant effects had statistical power of >0.999 at an effect size of >5% (OR > 1.05 or < 0.95).

**Table 1.** Logistic regression associations between deletion or duplications of tumour suppressor genes (TSG) and oncogenes (Onco) versus cancer incidence in the UKB cohort, excluding CNVs occurring in over 1000 participants. HE: hypothesized effect of the stated change, where ’+’ indicates a change expected to increase cancer risk while ’-’ indicates decreased cancer risk. Sig.: Statistical test significance codes (‘***’: p < 0.001, ‘**’:p < 0.01, ‘*’: p < 0.05, ‘.’: p < 0.10; NS: p > 0.10).

| Category | Extent | Type | Driver | HE | Est. | SE | z | p | N | OR | Sig. | Fits Hypothesis? |
| --- | --- | --- | --- | --- | --- | --- | --- | --- | --- | --- | --- | --- |
| TSG | Whole-Gene | Gain | Any | - | -0.007 | 0.049 | -0.145 | 0.885 | 2498 | 0.975 | NS | - |
| TSG | Whole-Gene | Gain | Driver | - | 0.040 | 0.051 | 0.778 | 0.436 | 2243 | 1.007 | NS | - |
| TSG | Whole-Gene | Loss | Any | + | 0.116 | 0.067 | 1.730 | 0.084 | 1256 | 1.076 | . | **Yes** |
| TSG | Whole-Gene | Loss | Driver | + | 0.146 | 0.069 | 2.133 | 0.033 | 1190 | 1.098 | * | **Yes** |
| TSG | Partial/CDS | Gain | Any | + | 0.001 | 0.036 | 0.037 | 0.971 | 4509 | 0.993 | NS | - |
| TSG | Partial/CDS | Gain | Driver | + | 0.032 | 0.034 | 0.952 | 0.341 | 5132 | 1.025 | NS | - |
| TSG | Partial/CDS | Loss | Any | + | -0.016 | 0.040 | -0.398 | 0.690 | 3735 | 0.984 | NS | - |
| TSG | Partial/CDS | Loss | Driver | - | 0.042 | 0.039 | 1.068 | 0.285 | 3777 | 1.027 | NS | - |
| Onco | Whole-Gene | Gain | Any | + | 0.012 | 0.050 | 0.248 | 0.804 | 2438 | 0.985 | NS | - |
| Onco | Whole-Gene | Gain | Driver | - | 0.011 | 0.054 | 0.199 | 0.842 | 2035 | 0.997 | NS | - |
| Onco | Whole-Gene | Loss | Any | - | 0.119 | 0.064 | 1.860 | 0.063 | 1388 | 1.068 | . | **Opposite** |
| Onco | Whole-Gene | Loss | Driver | + | 0.070 | 0.072 | 0.969 | 0.333 | 1117 | 1.043 | NS | - |
| Onco | Partial/CDS | Gain | Any | - | 0.015 | 0.032 | 0.455 | 0.649 | 5661 | 1.009 | NS | - |
| Onco | Partial/CDS | Gain | Driver | - | 0.017 | 0.040 | 0.432 | 0.666 | 3731 | 1.012 | NS | - |
| Onco | Partial/CDS | Loss | Any | - | 0.039 | 0.036 | 1.094 | 0.274 | 4650 | 1.023 | NS | - |
| Onco | Partial/CDS | Loss | Driver | - | 0.030 | 0.044 | 0.671 | 0.502 | 3044 | 1.015 | NS | - |

We then assessed the effect of CNV burden on cancer risk (Table 2), with the expectation that both increasing burden of all types (gains and losses; whole- and partial-gene) would contribute to increasing overall risk. We found that an increased burden of whole-gene losses significantly increased cancer risk (p = 0.00033, OR50: 1.050), as expected. Even explicitly excluding TSGs and oncogenes, this effect was still present (p = 0.00034, OR50: 1.050). Whole-gene gains were not found to have any statistically-significant effects on cancer risk, nor were gains or losses of partial-gene CDS.

**Table 2.** Effects of increasing CNV burden for each CNV type. OR50: Estimated odds-ratio for a participant carrying 50 CNVs of the specified type.

| Extent | Type | Driver | HE | Est. | SE | z | p | OR50 | Sig. | Supports |
| --- | --- | --- | --- | --- | --- | --- | --- | --- | --- | --- |
| Whole-Gene | Gain | All Genes | + | -4.58E-06 | 1.73E-04 | -0.027 | 0.979 | 1.000 | NS | - |
| Whole-Gene | Gain | Non-Driver | + | -3.37E-06 | 1.74E-04 | -0.019 | 0.985 | 1.000 | NS | - |
| Whole-Gene | Loss | All Genes | + | 9.71E-04 | 2.71E-04 | 3.584 | 3.38E-04 | 1.050 | *** | Yes |
| Whole-Gene | Loss | Non-Driver | + | 9.79E-04 | 2.73E-04 | 3.590 | 3.31E-04 | 1.050 | *** | Yes |
| Partial/CDS | Gain | All Genes | + | -8.00E-05 | 2.49E-04 | -0.321 | 0.748 | 0.996 | NS | - |
| Partial/CDS | Gain | Non-Driver | + | -7.97E-05 | 2.55E-04 | -0.313 | 0.754 | 0.996 | NS | - |
| Partial/CDS | Loss | All Genes | + | -5.89E-04 | 8.06E-04 | -0.730 | 0.465 | 0.971 | NS | - |
| Partial/CDS | Loss | Non-Driver | + | -6.10E-04 | 8.56E-04 | -0.713 | 0.476 | 0.970 | NS | - |

### Genome-wide copy number association tests between gene copy number and disease risk

Across all protein-coding genes, no significant associations were detected between gene duplications and cancer incidence. However, deletions of 34 genes were significantly associated with cancer incidence, after correction for multiple testing (*Figure 2A*; *Supplementary table 3*). Notably, 24 of these genes are located on chromosome 13, suggesting that the associations in this region may result from large CNVs encompassing multiple genes. Within this region, *KCNRG*, a potassium channel regulator and candidate tumour suppressor (Birerdinc et al., 2009; Cho et al., 2006), showed the strongest signal, alongside other cancer-related genes including *DLEU1*, *DLEU7*, *KPNA3*, and *TRIM13* (*Supplementary table 3*). Additional associations located on chromosomes 11, 14 and 22 were also grouped in clusters, indicating that their associations may also arise from large CNVs spanning multiple genes. On chromosome 14, a single gene, *KIAA0125*, was associated with cancer incidence. *KIAA0125* has been reported to act as a tumour suppressor (Hornung et al., 2018; Wang et al., 2021). Similarly, chromosome 20 contained one associated gene, *SNAI1*, a regulator of cancer progression (Fang et al., 2024; M. Li et al., 2024; Ryu et al., 2024). All associations indicated increased cancer risk with reduced gene copy number.

**Figure 2:**
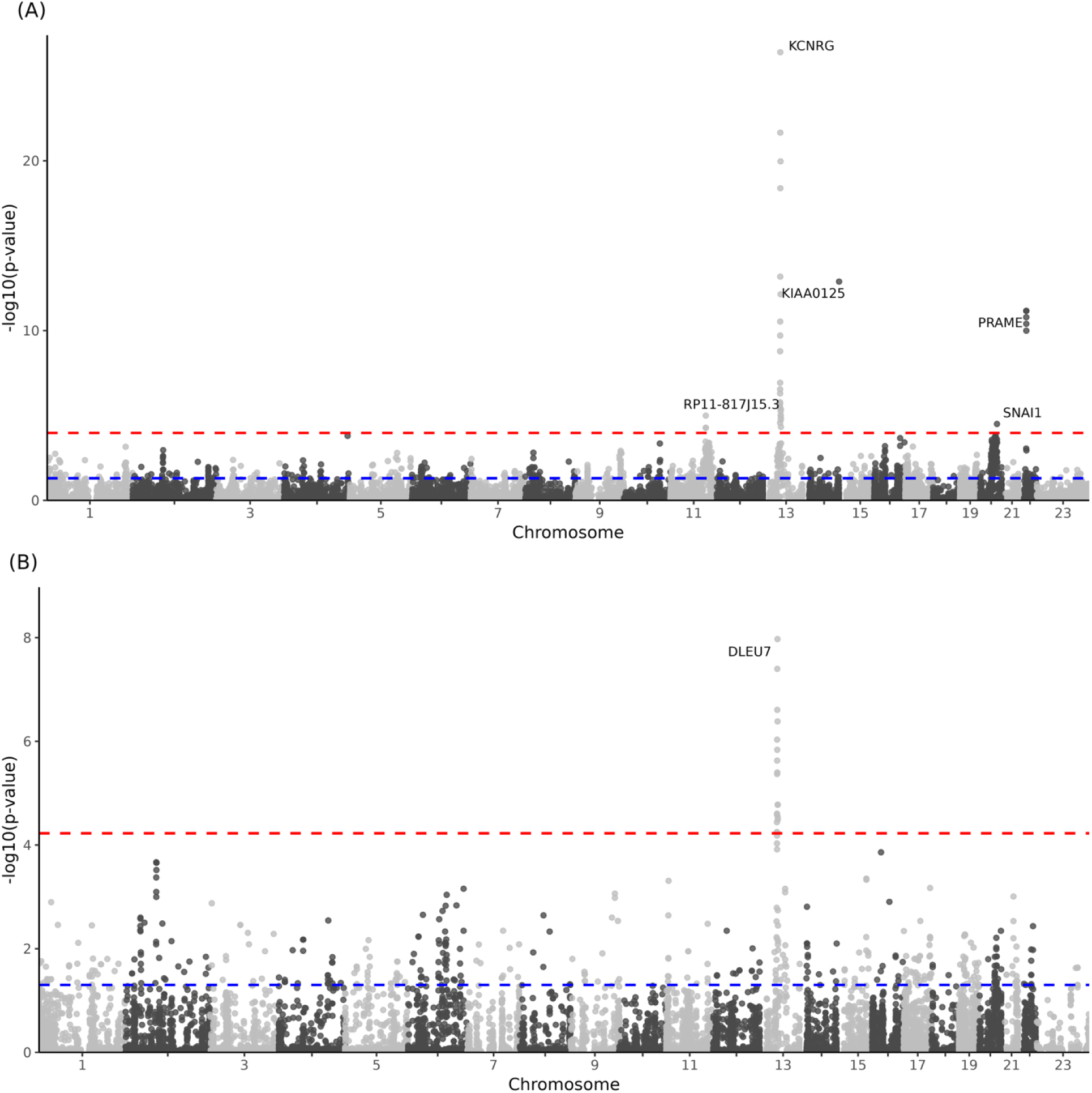
Manhattan plots showing association between the copy number of deleted genes and (**A**) incidence of any cancer type, and (**B**) skin cancer. Blue line represents p = 0.05 and red line represents adjusted p-value = 0.05.

We also performed separate analyses for skin, breast and prostate cancer, the three most common cancer types in the UK Biobank. No significant associations were identified for gene duplications in any of the cancer types. However, for skin cancer, we identified associations with deletions of 18 genes located on chromosome 13 (*Figure 2B*; *Supplementary table 3*). Among these, the gene with the strongest statistical significance was *DLEU7* (Deleted in Lymphocytic Leukaemia 7). No gene deletions were found to be significantly associated with either breast or prostate cancer.

We next extended the analysis to other diseases. Multiple associations were observed for both duplications (Supplementary *figure 2*; *Supplementary table 3*) and deletions (Supplementary *figure 3*; *Supplementary table 3*). Significant gene-level associations included: 48 duplications and 4 deletions with angina; 7 deletions with asthma; 21 duplications and 76 deletions with diabetes; 166 duplications with deep vein thrombosis (DVT); 14 duplications and 8 deletions with heart attack; and 18 duplications and 1 deletion with pulmonary embolism. No associations were found for stroke. The full list of associations can be found in Supplementary table 3. In all cases, copy number variation was associated with increased rather than reduced disease risk, suggestive of a causal association.

In addition to associations identified from whole-gene deletions, we also observed associations between deletions that disrupt the coding sequences of protein-coding genes and disease phenotypes. These included deletions affecting *REV3L*, the catalytic subunit of DNA polymerase, which were associated with skin cancer (Supplementary figure 4B), and deletions affecting *SNRNP27*, a small nuclear ribonucleoprotein, which were associated with breast cancer (Supplementary figure 4C). Additional associations were observed between partial gene deletions and disease phenotypes (Supplementary figure 5; Supplementary data 4).

To compare the performance of our gene-centric framework, we conducted a standard CNV association test using two distinct methods of CNV aggregation: a density-based procedure (Kim et al., 2012) and a reciprocal overlap approach (da Silva et al., 2019). In contrast to our primary gene-centric analysis, neither approach yielded any deletions that were significantly associated with cancer incidence after multiple test correction (FDR < 0.05*; Supplementary table 4*). This was consistent across all tested parameters for both aggregation methods.

## Discussion

Dosage sensitivity has been proposed as a key mechanism underlying the pathogenicity of CNVs (Rice & McLysaght, 2017), but traditional CNV studies often analyse broad genomic regions, making it difficult to resolve causal genes. Here, we applied a gene-centric framework to assess the effects of whole-gene copy number changes on gene expression and disease risk.

Relatively few genes showed variable copy number across individuals, likely reflecting the fact that variable copy number is itself uncommon for most genes. Many genes are likely to be under purifying selection to maintain their expression levels within a narrow range, reducing the chances of observing significant expression-copy number correlations in a healthy cohort (Makino & McLysaght, 2010; Papp et al., 2003; Rice & McLysaght, 2017). Among genes with detectable copy number variation, most exhibited the expected positive relationship between copy number and expression, consistent with dosage sensitivity and prior work (Collins et al., 2022; Yamasaki et al., 2020). Immune genes were enriched among genes showing such correlations, which is likely driven by the fact that CNVs are enriched in regions encoding immune genes (Nguyen et al., 2006), reflecting the adaptive role of structural variation in generating immunological diversity (Nguyen et al., 2006; Schaschl et al., 2009).

A small number of genes had a negative correlation between copy number and gene expression. Previous studies of CNVs in mammalian cells have shown similar results, with 30% of genes showing a decreased expression with increased copy number (R. T. Wang et al., 2011). Various explanations have been proposed for these cases, including regulatory feedback loops where overexpression leads to the activation of inhibitory signals (Henrichsen et al., 2009; Schlattl et al., 2011). However, in our own data, seven genes exhibited unexpected negative correlations, three of which were HLA pseudogenes. We propose this may be the result of spurious CNV calls due to the highly polymorphic nature of the HLA region of the genome, as reported in previous studies (Fawcett et al., 2022).

To validate our assumption that additional gene copies increase the opportunity for mutation, we compared somatic mutation rates between X-linked genes in males and females. Genes with an extra copy in females exhibited approximately twice as many mutations, whereas pseudoautosomal genes showed no sex difference. Interestingly, the effect is much stronger in inactivated genes, which could be explained by transcription-coupled repair (reviewed in (Hanawalt & Spivak, 2008), given that genes escaping X inactivation remain transcriptionally active on both chromosomes, thus leading to a lower mutation rate.

Despite most known changes in gene copy number being deleterious, there are a few examples where it may have protective effects (Gonzalez et al., 2005; Greenhill, 2014; León-Mimila et al., 2018). Based on the two-hit model of tumorigenesis (Knudson, 1971), we hypothesised that additional tumour suppressor gene (TSG) copies might be protective, while deletions could elevate risk. Some of these expected patterns were observed, with whole-gene losses of TSGs associated with higher cancer risk. Oncogenes were expected to show the inverse of this pattern, but they instead demonstrated the same effects (albeit at marginal significance), with whole-gene losses increasing cancer risk This was at odds with our increased oncogenic target-area hypothesis. We then considered the effects of CNV burden overall, analysing all whole- and partial-gene gains and losses across the genome. We expected that participants enriched for CNVs in general would experience greater lifetime cancer risk. This effect was observed for whole-gene losses. Furthermore, even when all cancer-associated genes (both oncogenes and TSGs) were excluded, a highly significant effect was still present. This suggests that, while some key genes do have strong associations to cancer, genomic dysregulation broadly may itself be a cancer risk factor.

To move beyond known cancer genes and leverage the power of our gene-centric approach, we expanded our analysis to a genome-wide association study between the copy number of all protein-coding genes and cancer incidence. Deletions in the 13q14 region were associated with increased overall cancer incidence and increased incidence of skin cancer. While associations between somatic alterations in the 13q14 region and cancer are widely established (Birerdinc et al., 2009; Khalid et al., 2021; Palamarchuk et al., 2010), reports of germline risk factors in this locus are less common. Although germline mutations in *RB1* (a gene within 13q14) defines a predisposition to cancer (Knudson, 1971; Rojanaporn et al., 2018), this gene falls outside of our associated locus. Therefore, our findings point to a separate, independent germline risk factor within this cytogenetic band. Here, we identify *KCNRG*, a putative tumour suppressor (Ivanov et al., 2003), as the gene with the strongest association with overall cancer incidence.

Our ability to detect novel associations between gene copy number and phenotypes is underscored by a direct comparison with standard CNV-association methods. When we aggregated CNVs into broader genomic regions using density-based and reciprocal overlap approaches, none of the resulting regions were significantly associated with cancer incidence after multiple testing correction. The failure of these conventional methods to detect the signal captured by our gene-centric approach suggests that aggregating CNVs into large, heterogeneous regions can dilute statistical power. This contrast highlights the advantage of a gene-focused strategy for pinpointing discrete, biologically relevant risk loci that might otherwise be obscured.

Additional associations between deletion and cancer incidence include *KIAA0125*, a long non-coding RNA. *KIAA0125* has previously been identified as a candidate tumour suppressor with prior studies implicating its dysregulation in cancer progression (Diniz et al., 2019; Yang et al., 2019). However, to the best of our knowledge, this is the first evidence directly associating *KIAA0125* deletion with cancer, supporting its proposed tumour suppressive role.

Beyond cancer, we identified potentially novel associations between gene copy number and other disease phenotypes. For example, potentially novel associations include deletions of *CHRNA7* and *P2RX1* in relation to asthma. While both genes have been implicated in immune regulation and inflammation (Bencherif et al., 2011; Thompson et al., 2021; Wongtrakool et al., 2012), there is no prior evidence linking their deletions to asthma. Similarly, we observed associations between deletions of *TLR5* and *SMYD4* and heart attack; however, although these genes have previously been implicated in cardiovascular health (Parapanov et al., 2015; Rueda-Robles et al., 2021; Xiao et al., 2018), we found no evidence that their deletions are linked to heart attack incidence. Additionally, we found duplications of *HIST1H2BM, HIST1H4J, CAPN3,* and *ZNF157* to be linked to an increased risk of heart attack, none of which have been previously associated with cardiovascular risk. A full list of associations is provided in the Supplementary table 3.

In conclusion, we have explored the impact of gene copy number gains and losses on gene expression and disease risk using data from the GTEx consortium and the UK Biobank. We found many genes exhibited the expected positive relationships between copy number and gene expression; however, we also identified cases of negative correlation, potentially explained by technical artifacts or regulatory feedback mechanisms. The hypothesized relationships between gains and losses of TSGs and oncogenes were broadly not supported, with the sole exception of TSG loss associated with increased cancer risk. Instead, whole-gene copy-loss burden broadly was strongly associated with increased cancer risk, while no significant associations were found for CDS events or copy-gains. Finally, we identified other gene copy number variations associated with disease phenotypes, with all identified associations being pathogenic rather than protective. This research contributes to our understanding of gene dosage effects and the broader phenotypic consequences of variations in gene copy number, highlighting associations between individual genes and disease phenotypes, including implications for cancer susceptibility.

## Data Availability

This study used data from the GTEx consortium and UK Biobank. Results generated from this study are included in the manuscript and supplementary material. The code used to generate results presented in this study are available from our github: https://github.com/sophie-03/gene_centric_CNV_analysis

## Acknowledgements

This research was funded by Research Ireland through the Research Ireland Centre for Research Training in Genomics Data Science under grant number 18/CRT/6214 and has been conducted using the UK Biobank Resource under Application Number 74519. UK Biobank has ethical approval from the North West Multi-centre Research Ethics Committee (MREC), and all participants provided informed consent. We would like to thank members of the Seoighe lab: Harrison Anthony, Tyler Medina and Mehak Chopra, at the University of Galway for valuable discussions regarding this research.

---

**Supplementary figure 1:**
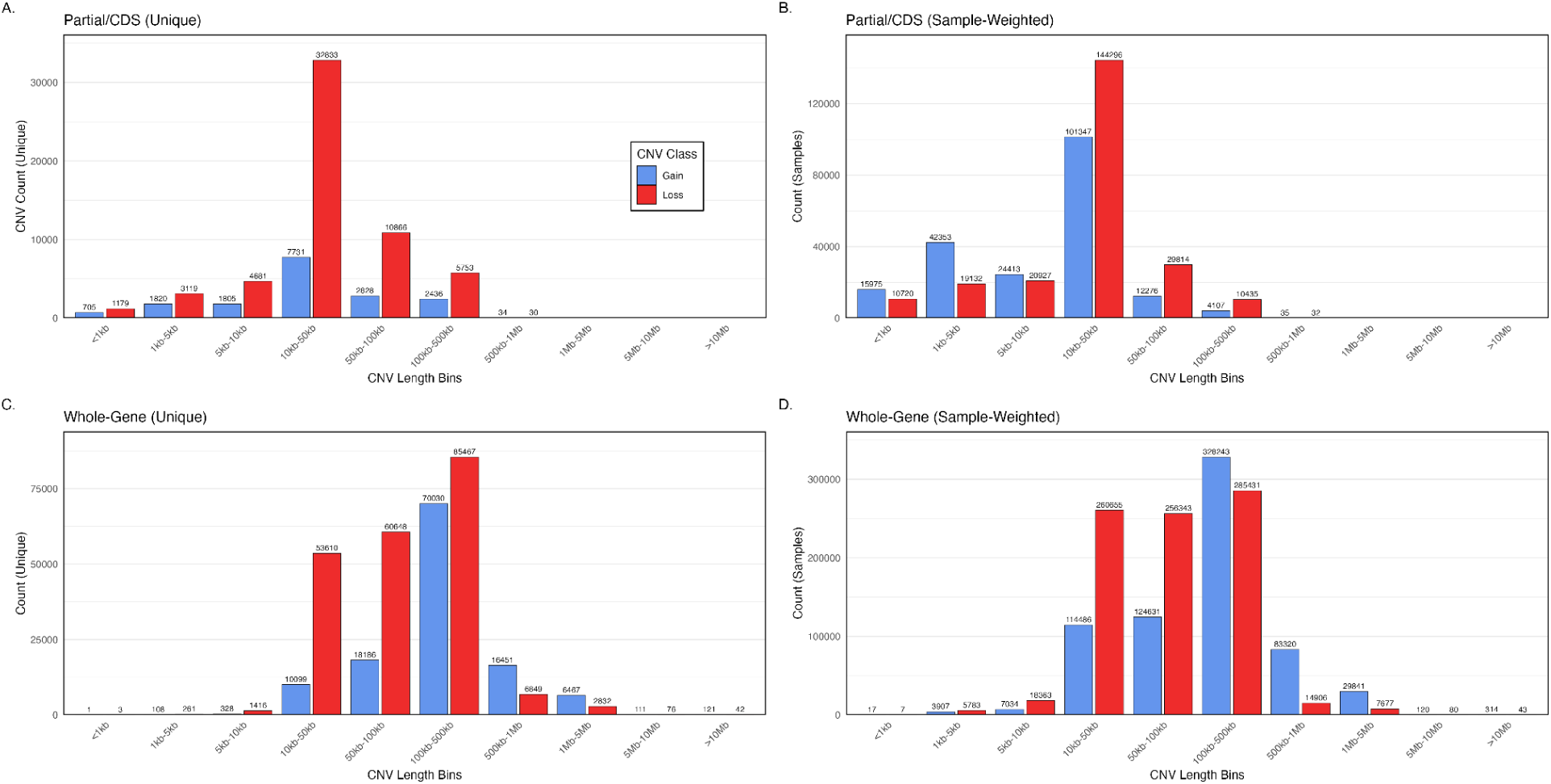
Distribution of lengths of all CNV events included in study. **(A)** CDS-impacting partial-gene CNVs as counts of unique breakpoint sets. **(B)** CDS-impacting partial-gene CNVs weighted by occurrences in samples. **(C)** Whole-gene CNVs as counts of unique breakpoint sets. **(D)** Whole-gene CNVs weighted by occurrences in samples. Gains are shown in blue and losses in red for all subfigures.

**Supplementary figure 2:**
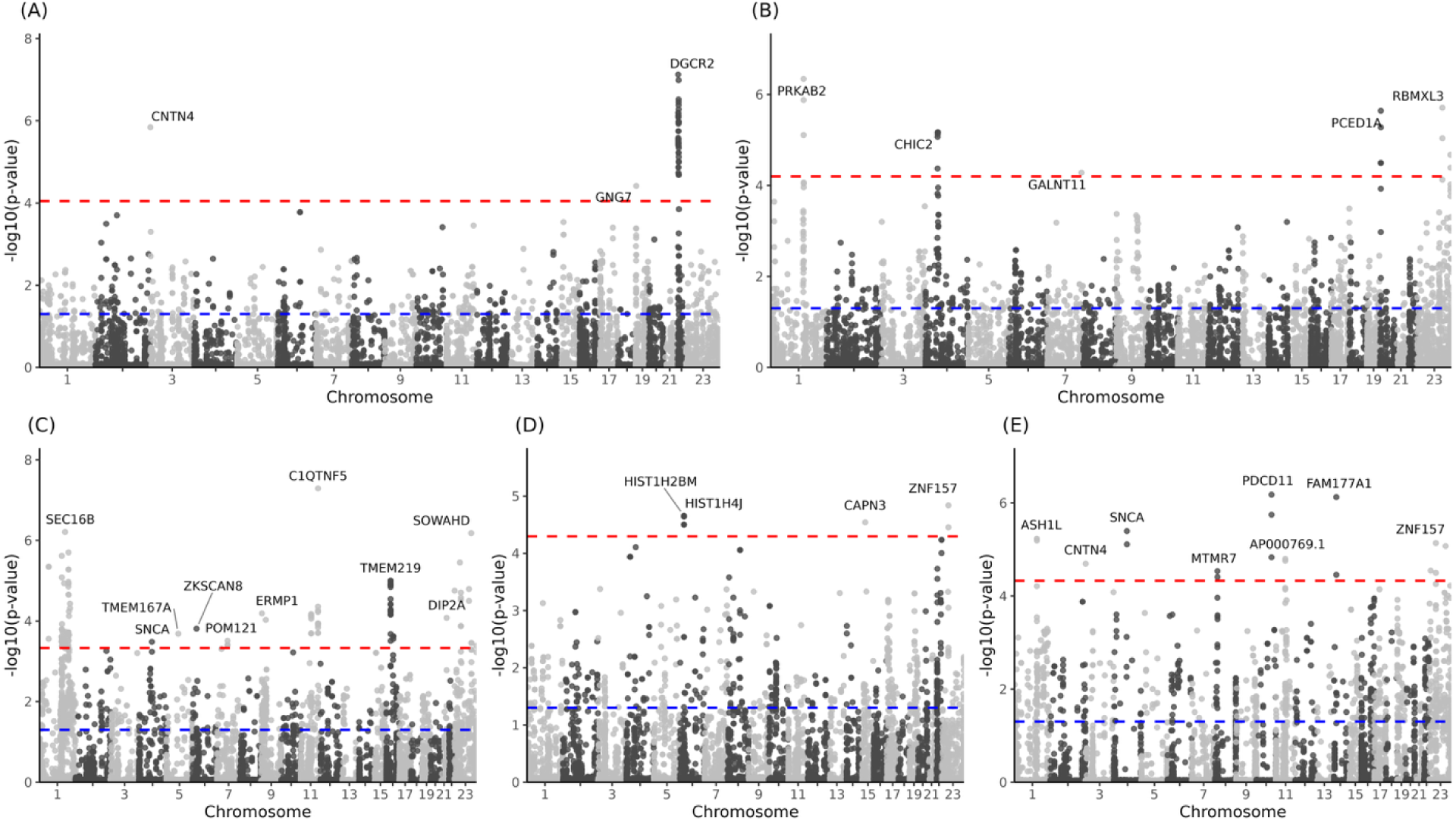
Manhattan plots showing association between the copy number of duplicated genes and disease phenotypes: (**A**) angina, (**B**) diabetes, (**C**) DVT, (**D**) heart attack and (**E**) pulmonary embolism. Blue line represents p = 0.05 and red line represents adjusted p-value = 0.05.

**Supplementary figure 3:**
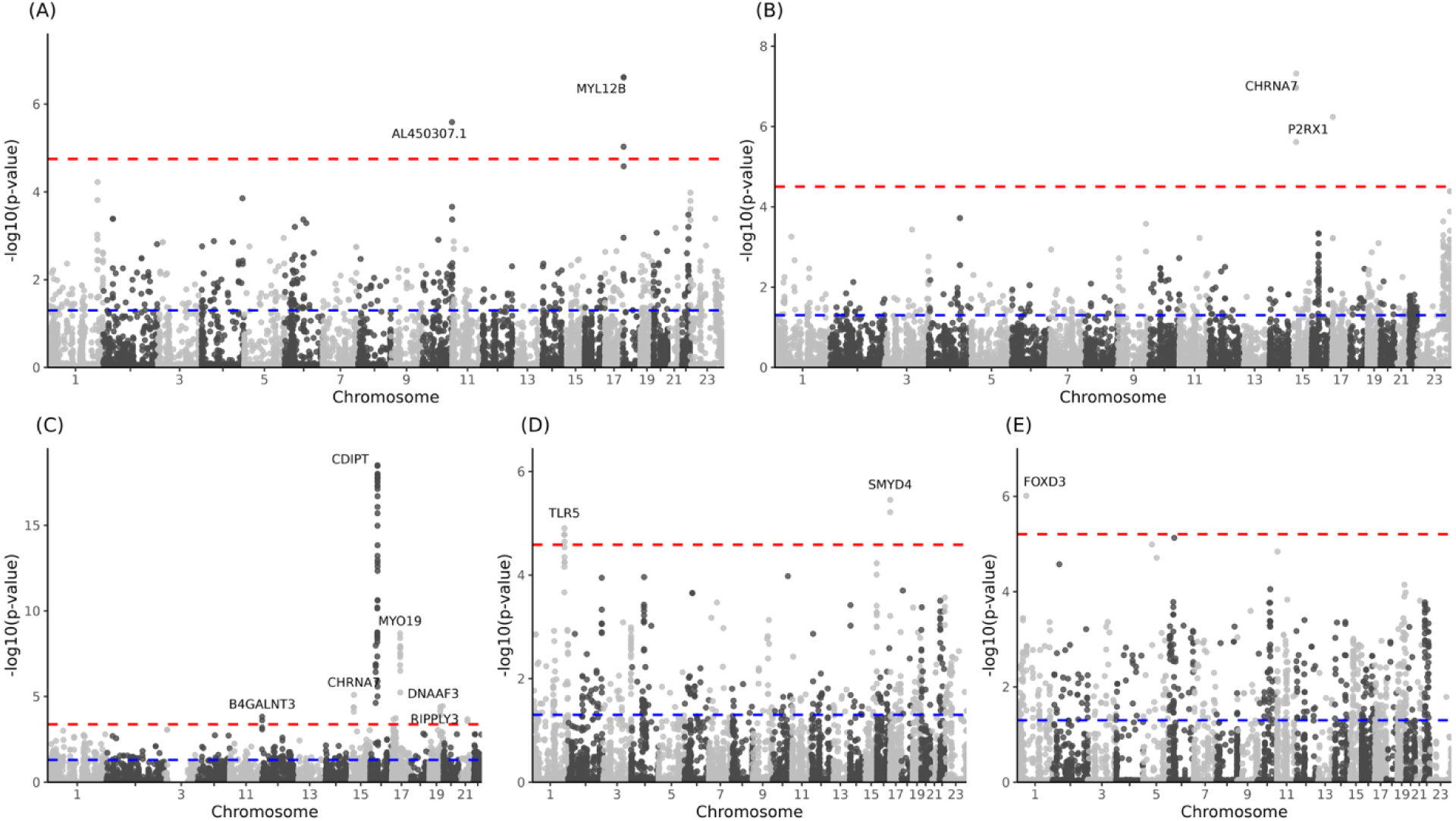
Manhattan plots showing association between the copy number of deleted genes and disease phenotypes: (**A**) angina, (**B**) asthma, (**C**) diabetes, (**D**) heart attack and (**E**) pulmonary embolism. Blue line represents p = 0.05 and red line represents adjusted p-value = 0.05.

**Supplementary figure 4:**
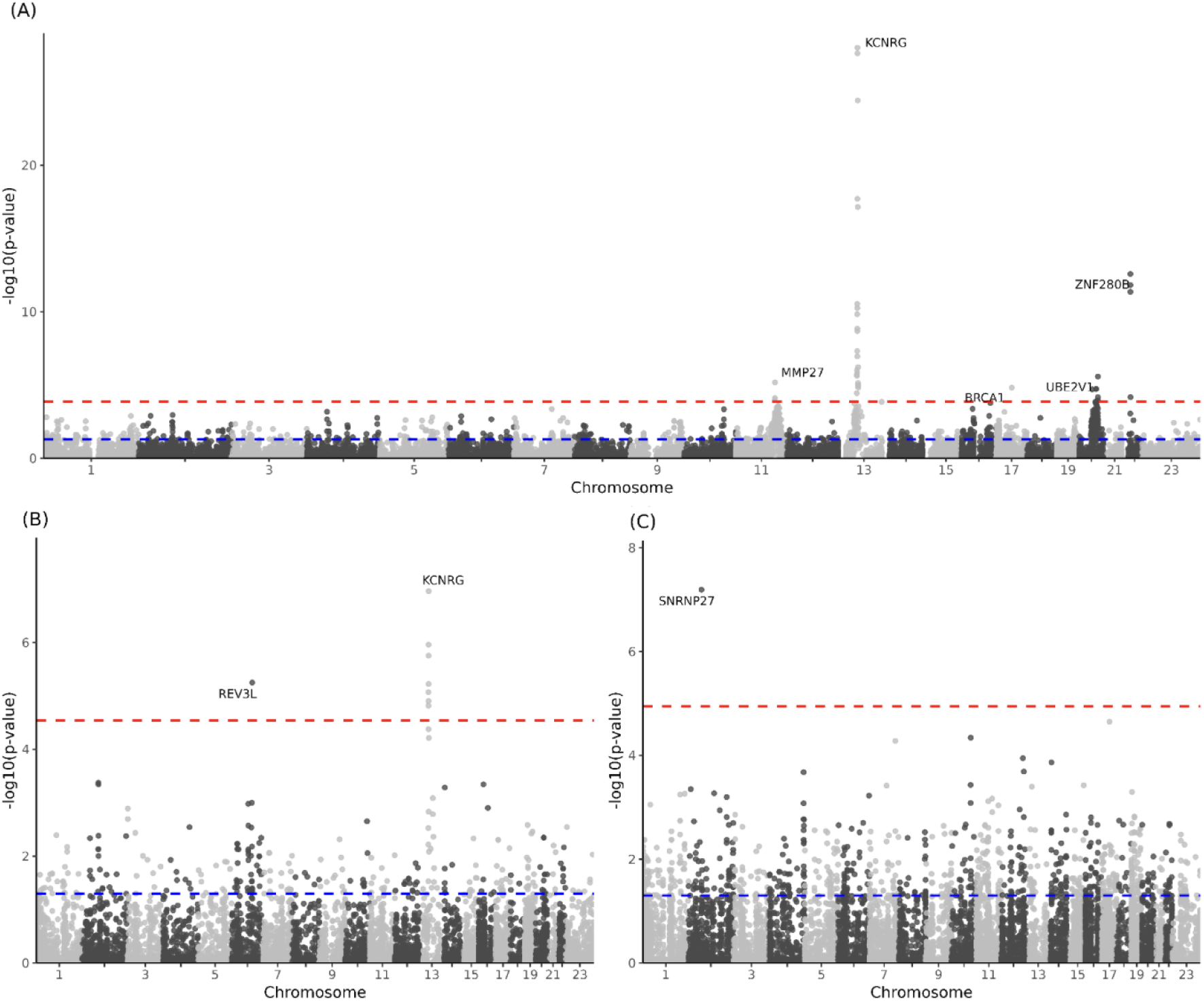
Manhattan plots showing association between deletions disrupting protein-coding genes and (**A**) incidence of any cancer type, (**B**) skin cancer and (**C**) breast cancer. Blue line represents p = 0.05 and red line represents adjusted p-value = 0.05.

**Supplementary figure 5:**
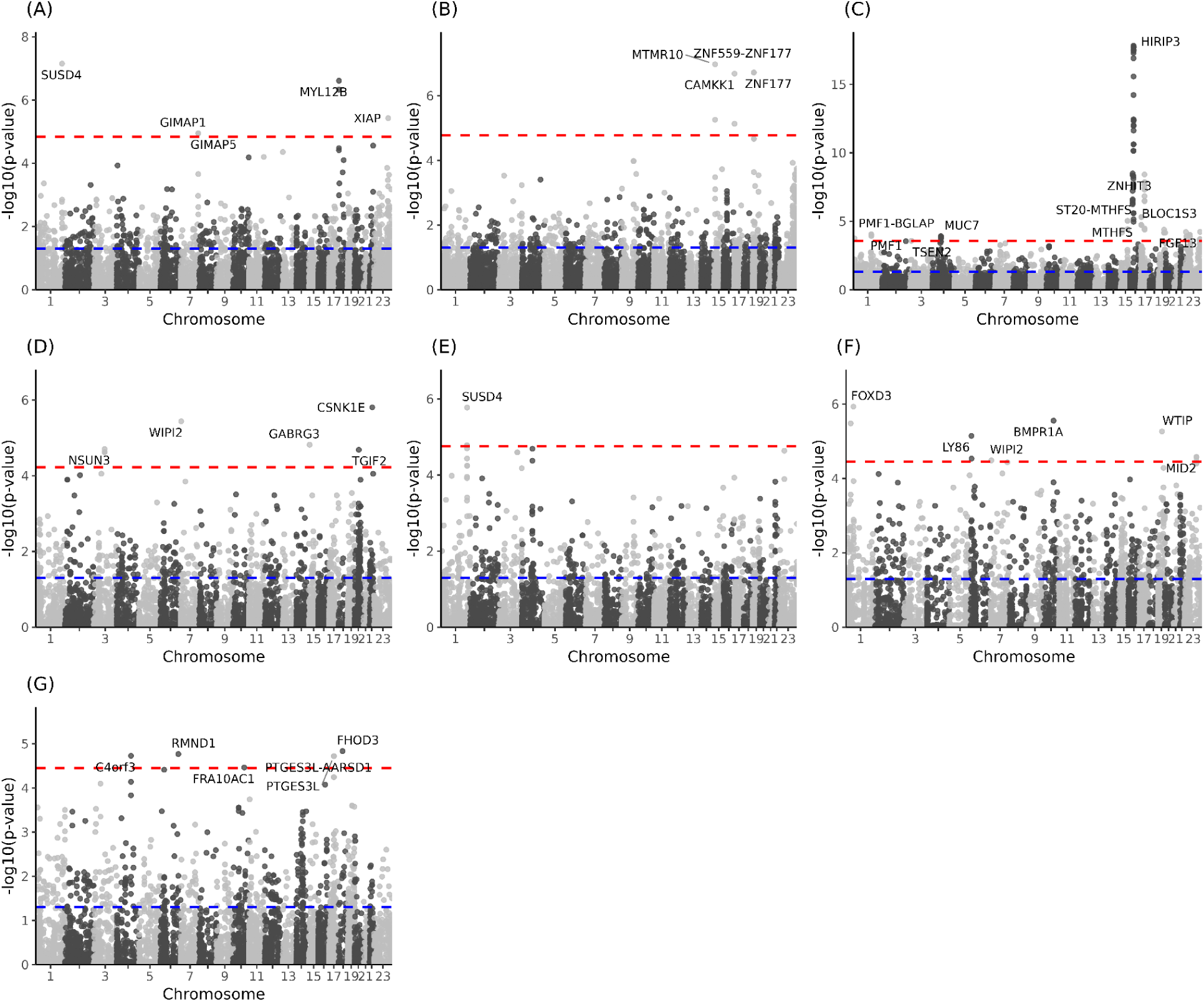
Manhattan plots showing association between deletions disrupting protein-coding genes and (**A**) angina, (**B**) asthma, (**C**) diabetes, (**D**) deep vein thrombosis, (**E**) heart attack, (**F**) pulmonary embolism and (**G**) stroke. Blue line represents p = 0.05 and red line represents adjusted p-value = 0.05.

## Notes

### Competing Interest Statement

The authors have declared no competing interest.

